# A novel epigenetic clock for rhesus macaques unveils an association between early life adversity and epigenetic age acceleration

**DOI:** 10.1101/2024.10.08.617208

**Authors:** Gabriel Bronk, Roy Lardenoije, Laura Koolman, Claudia Klengel, Shu Dan, Brittany R. Howell, Elyse L. Morin, Jerrold S. Meyer, Mark E. Wilson, Kelly F. Ethun, Maria C. Alvarado, Jessica Raper, Hector Bravo-Rivera, Margaux M. Kenwood, Patrick H. Roseboom, Gregory J. Quirk, Ned H. Kalin, Elisabeth B. Binder, Mar M. Sanchez, Torsten Klengel

## Abstract

Because DNA methylation changes reliably with age, machine learning models called epigenetic clocks can estimate an individual’s age based on their DNA methylation profile. This epigenetic measure of age can deviate from one’s true age, and the difference between the epigenetic age and true age, known as epigenetic age acceleration (EAA), has been found to directly correlate with morbidity and mortality in adults. Emerging evidence suggests that EAA is also associated with aberrant health outcomes in children, making epigenetic clocks useful tools for studying aging and development. We developed two highly accurate epigenetic clocks for the rhesus macaque, utilizing 1,008 blood samples from 690 macaques between 2 days and 23.4 years of age with diverse genetic backgrounds and exposure to environmental conditions. The first clock, which is trained on all samples, achieves a Pearson correlation between true age and predicted age of 0.983 and median absolute error of 0.210 years. To study phenotypes during development, the second clock is optimized for macaques younger than 6 years and achieves a Pearson correlation of 0.974 and a median absolute error of 0.148 years. Using the latter clock, we investigated whether epigenetic aging is affected by early life adversity in the form of infant maltreatment. Our data suggests that maltreatment and increased hair cortisol levels are associated with epigenetic age acceleration right after the period of maltreatment.

## Introduction

As an individual ages, the probability of DNA methylation at specific sites across the genome monotonically increases or decreases. Prior studies have taken advantage of the association between DNA methylation and age by creating machine learning models called epigenetic age estimators or epigenetic clocks that predict age based on DNA methylation profiles in a variety of human tissues. Among the most well-known estimators are clocks developed by Hannum et al.^1^ and Horvath^2^ that linearly regress age on DNA methylation. Moreover, prior studies investigating differences between true age (referred to as “chronological age”) and the age predicted by epigenetic clocks (a measure of “biological age”) have suggested that some individuals appear biologically older or younger than their chronological age. Differences between predicted age and true age, termed age acceleration or age deceleration, offer an opportunity to better understand factors influencing aging processes.

Studies using the aforementioned Horvath^2^ or Hannum^1^ human clocks have discovered associations between epigenetic age acceleration and health in both adults and children. In adults, age acceleration predicts risk of all-cause mortality^3,4^ and mortality due to cancer^3^ and is directly correlated with a decline in cognitive abilities^5,6^ and physical fitness.^5^ In children, studies have found that age acceleration directly correlates with early onset of puberty^7^ and risk of asthma and allergies^8^ and inversely correlates with hippocampal volume.^9^ These studies calculated age acceleration using samples of peripheral blood,^3–5,7,8^ saliva,^9^ or the dorsolateral prefrontal cortex.^6^ Epigenetic age acceleration has been found to be heritable and influenced by environmental factors.^2^ Therefore, epigenetic clocks may be useful tools for discovering genetic or environmental factors that speed up or slow down the aging process.

Trauma exposure is one such factor that has been suggested to cause epigenetic age acceleration. Using Horvath’s clock,^2^ several studies uncovered an association with trauma exposure: Boks et al. found that soldiers who experienced a large number of traumatic events during a war had statistically significant epigenetic age acceleration 6 months after their tour of duty compared to soldiers who experienced few traumatic events.^10^ Jovanovic et al. performed a cross-sectional study in children ages 6 to 13 years old and found that age acceleration is directly correlated with the frequency at which the children have experienced violence.^11^ Sumner et al. performed a cross-sectional study in children ages 8 to 16 years old, finding age acceleration is directly correlated with an early life adversity (ELA) score, which includes multiple types of traumatic events such as the number of experiences of physical, sexual and emotional abuse throughout an individual’s childhood.^12^ These studies suggest that childhood and adult trauma exposure is associated with age acceleration, which in turn may be related to the broad effects of trauma on general morbidity and mortality. However, human studies may be confounded by other environmental factors and may not provide evidence for causal relationships between trauma exposure and age acceleration.

Rodents and non-human primates are viable translational research models for studying genetic and environmental factors influencing aging, including ELA. Rhesus macaques experience aging processes similar to humans and offer unique translational value due to their close relationship with humans with respect to genetic, neural, behavioral, endocrine, and immune system phenotypes.^13^ In addition, both in the wild and in captivity some rhesus macaques experience ELA in the form of infant maltreatment.^14–17^ Two epigenetic clocks for rhesus macaque have been published,^18,19^ but to the best of our knowledge, no association between age acceleration and biological or environmental factors has been found. This is in contrast to a study in baboons where an association between social hierarchy and epigenetic aging was established.^20^ Here we develop two precise epigenetic age estimators from a large, diverse cohort of rhesus macaques between 2 days and 23.4 years of age and provide evidence for the effects of early life adversity on age acceleration in rhesus macaques.

## Results

### Two Epigenetic Clock Models Accurately Predict Chronological Age in Rhesus Macaques

We investigated DNA methylation profiles in 1,008 peripheral blood samples from rhesus macaques (*Macaca mulatta*) between 2 days and 23.4 years of age (mean = 2.5 years, standard deviation = 3.4 years). Samples were collected from a total of 690 macaques, with blood drawn at multiple time points in 86 of the individuals. Samples were obtained predominantly from young animals with 85% of samples from animals between 0 to 3 years of age. The distributions of ages and rhesus macaque cohorts in our dataset are depicted in **Table S1**. Epigenetic clocks were trained on animals from the Wisconsin National Primate Research Center (WNPRC), the Emory National Primate Research Center (ENPRC), and the Caribbean Primate Research Center (CPRC), utilizing five separate cohorts with varying ages, genetic backgrounds and exposure to environmental conditions, ensuring that the clocks are generalizable to other rhesus macaque populations.

The epigenetic clock is a multiple linear model, where the dependent variable is the animal’s age, and the independent variables are DNA methylation levels at CpG sites throughout the genome. To achieve maximal model accuracy, we used only reliable CpG probes, determined through intraclass correlation coefficients for each CpG in replicate samples (see *Methods, DNA Methylation Data Processing*).^21^ Our first model was trained on all 1,008 samples and fit using elastic net regularization, which selected 344 CpG sites (**Table S2**). We name this model the *Epigenetic Clock for Rhesus Macaques* (ECRM) to differentiate it from an additional clock developed for young individuals (discussed below). Using nested cross validation, we computed the following performance metrics for ECRM: the Pearson correlation between true age and predicted age is 0.983, the median absolute error is 0.210 years, the mean absolute error is 0.365 years, and the root mean square deviation (RMSD) is 0.640 years. **Figure 1** displays the true ages and predicted ages for all samples. The ECRM model is highly accurate for rhesus macaques, especially in the first 15 years of life.

**Figure 1:**
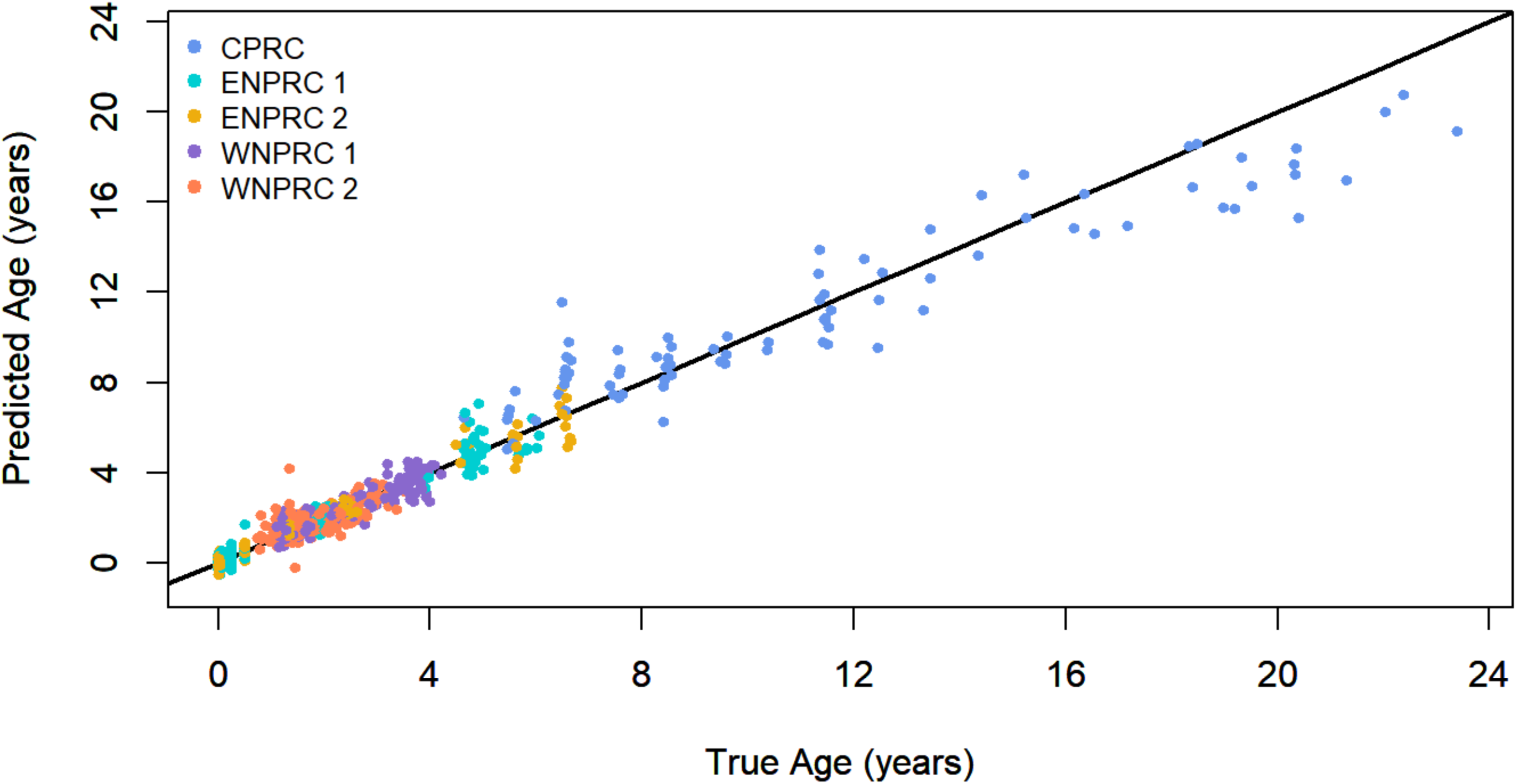
Comparison of true ages to ages predicted by the *Epigenetic Clock for Rhesus Macaques* (ECRM). Each colored dot represents a measurement from an individual, and the color represents the cohort in which the individual was raised. The black line is a line of slope 1 through the origin. Ages are predicted for all 1,008 rhesus macaque samples using 10-fold nested cross validation. CPRC: Caribbean Primate Research Center; ENPRC: Emory National Primate Research Center; WNPRC: Wisconsin National Primate Research Center).

To better study aging and developmental processes in young rhesus macaques, we also created a version of the epigenetic clock that is even more accurate than ECRM during the developmental period, which is between birth and 6 or 7 years of age (6 years for rhesus macaques is equivalent to about 24 years for humans).^22–24^ We name it the *Epigenetic Clock for Young Rhesus Macaques* (ECYRM). This model uses 453 CpG sites (**Table S3**), selected by elastic net regularization, and **Figure 2** displays the true ages and predicted ages for all samples in ECYRM. For individuals below the age of 6 years, ECYRM achieves a Pearson correlation between true age and predicted age of 0.974, a median absolute error of 0.148 years, a mean absolute error of 0.208 years, and an RMSD of 0.299 years. For comparison, for animals below the age of 6 years, ECRM achieves a Pearson correlation between true age and predicted age of 0.958, a median absolute error of 0.195 years, a mean absolute error of 0.269 years, and an RMSD of 0.388 years. This improvement of ECYRM compared to ECRM for the first 6 years is statistically significant as the Pearson correlations (and 95% CIs) for ECYRM and ECRM are 0.974 (0.971-0.977) and 0.958 (0.953-0.963), respectively. Such accuracy during the developmental period has not been demonstrated in prior studies of rhesus macaques.

**Figure 2:**
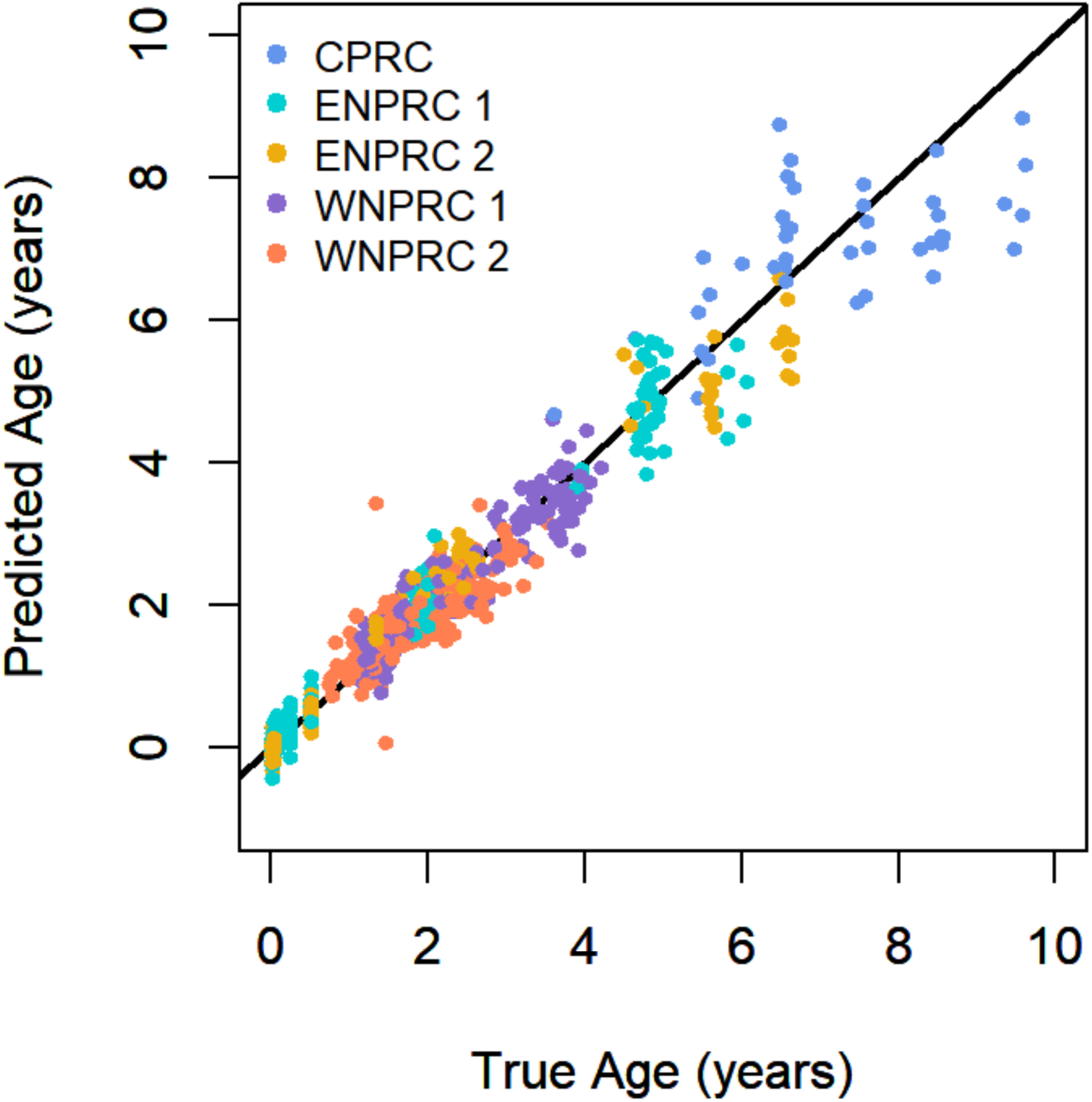
Comparison of true ages to ages predicted by the *Epigenetic Clock for Young Rhesus Macaques* (ECYRM). Each colored dot represents a measurement from an individual, and the color represents the cohort in which the individual was raised. The black line is a line of slope 1 through the origin. Ages are predicted for the 963 rhesus macaque samples under the age of 10 years using 10-fold nested cross validation. The epigenetic clock is most accurate for individuals under the age of 6 years. CPRC: Caribbean Primate Research Center; ENPRC: Emory National Primate Research Center; WNPRC: Wisconsin National Primate Research Center).

ECYRM was trained in the same manner as ECRM except it is only trained on samples from macaques under the age of 10 years (n = 963) (see *Methods* for the rationale for an age range of 0-10 years).

### ELA in Form of Infant Maltreatment and Hair Cortisol Levels in Rhesus Macaques Are Associated with Age Acceleration

Prior studies in human clinical samples have provided evidence for the association of ELA with epigenetic age acceleration.^25^ To test whether epigenetic age acceleration is associated with ELA in rhesus macaques, we utilized blood DNA methylation data from developing rhesus macaques ages 2 days to 2.2 years from a translational longitudinal study on infant maltreatment in which rhesus macaque mothers abused and neglected their infants,^14–17^ which we refer to as the ENPRC 1 cohort (see **Table S1** and *Methods, Cohort Information* for details).

Prior to predicting epigenetic age in the ENPRC 1 cohort, we retrained the ECYRM model, excluding ENPRC 1 individuals from the training dataset to prevent data leakage, resulting in a training set of 771 samples below 10 years of age. We then used this retrained clock (which we refer to as ECYRM_modified_) to accurately predict epigenetic ages of the ENPRC 1 macaques (Pearson correlation between true age and predicted age was 0.975, median absolute error was 0.146 years, mean absolute error was 0.295 years, and RMSD was 0.487 years). Finally, we calculated epigenetic age accelerations by subtracting chronological age from predicted age. One outlier sample was removed from the control group due to its large inconsistency with the age accelerations of the rest of the control group (see *Methods, Age Acceleration Outlier Detection*).

To quantify the effect of infant maltreatment on epigenetic age acceleration, we performed Bayesian multiple linear regression with epigenetic age acceleration as the dependent variable, and maltreatment and the macaque’s sex as the independent variables (the maltreatment variable is designated as 0 for the control group and 1 for the maltreatment group). The multiple linear model and Bayesian parameter estimation are described further in *Methods, Modeling Age Acceleration*.

As blood samples were drawn at multiple time points, we were able to monitor epigenetic age acceleration before maltreatment (i.e. at postnatal day 2), during maltreatment (age 2 weeks to 6 months), and well after maltreatment (at 1.8-2.2 years old). Multiple linear regression models were fit separately for each time point, and **Table 1** shows the age acceleration due to maltreatment as estimated by the regressions. Between 2 days and 3 months of age, no age acceleration difference between the maltreatment and control group was detected. Specifically, we found that the 95% confidence intervals (CIs) of age acceleration include 0 years, suggesting insufficient evidence to conclude that maltreatment affects age acceleration (the 95% CI is technically the credible interval, which is the Bayesian equivalent of the frequentist 95% confidence interval). However, at 6 months of age, we found that the epigenetic age acceleration in maltreated individuals is 0.15 years (95% CI: 0.0064 to 0.29) greater than for control individuals. We also investigated whether infant maltreatment has lasting effects on age acceleration beyond the period of maltreatment. For individuals between ages 1.8 and 2.2 years, we did not find an age acceleration associated with their prior maltreatment during infancy (**Table 1**).

**Table 1.** Age accelerations associated with maltreatment. Epigenetic age acceleration associated with maltreatment was calculated at six different time points between birth and 2.2 years. 95% CIs suggest that infant maltreatment is associated with a positive age acceleration at 6 months of age but not before (day 2), during (week 2, month 1, month 3) or long after maltreatment (2 years).

| Age | Age (years) | Age Acceleration (years) [95% CI] | Sample Size (Number in maltreatment group, number in control group) |
| --- | --- | --- | --- |
| 2 days | 0.0055 | 0.041 [-0.080 to 0.16] | 25 (14, 11) |
| 2 weeks | 0.038 | -0.0016 [-0.15 to 0.15] | 17 (8, 9) |
| 1 month | 0.083 | 0.15 [-0.0059 to 0.31] | 22 (15, 7) |
| 3 months | 0.25 | -0.069 [-0.15 to 0.017] | 31 (16, 15) |
| 6 months | **0.5** | **0.15 [0.0064 to 0.29]** | **23 (13, 10)** |
| 21 - 27 months | 1.8-2.2 | -0.023 [-0.26 to 0.22] | 30 (14, 16) |

To further quantify the relationship between maltreatment and age acceleration beyond a categorical approach, we analyzed the available hair cortisol samples from the 6-month-old macaques (n = 21 – i.e. 11 maltreated and 10 control macaques). Hair cortisol provides a more precise measure of chronic stress exposure due to cortisol accumulation in hair over time. Previously, the research group at the ENPRC has reported higher hair cortisol accumulation in 6-month maltreated infants than in controls.^17^ We performed multiple linear regression with age acceleration as the dependent variable, and hair cortisol and sex as the independent variables. Through Bayesian parameter estimation, we found that for animals with a chronological age of 0.5 years, the effect size of hair cortisol for age acceleration is 0.10 years (95% CI of 0.027 to 0.17 years), indicating that for a one standard deviation increase in hair cortisol levels, there is a corresponding increase in age acceleration of approximately 0.10 years. Animals that differ in hair cortisol by two standard deviations differ in age acceleration by 0.20 years (95% CI of 0.054 to 0.34 years). The 95% CI does not overlap with 0 years, indicating that there is a positive association between hair cortisol and age acceleration (in Bayesian statistics, this is equivalent to finding a significant p-value in frequentist statistics). **Figure 3** displays the age accelerations vs. hair cortisol levels at 6 months.

**Figure 3.**
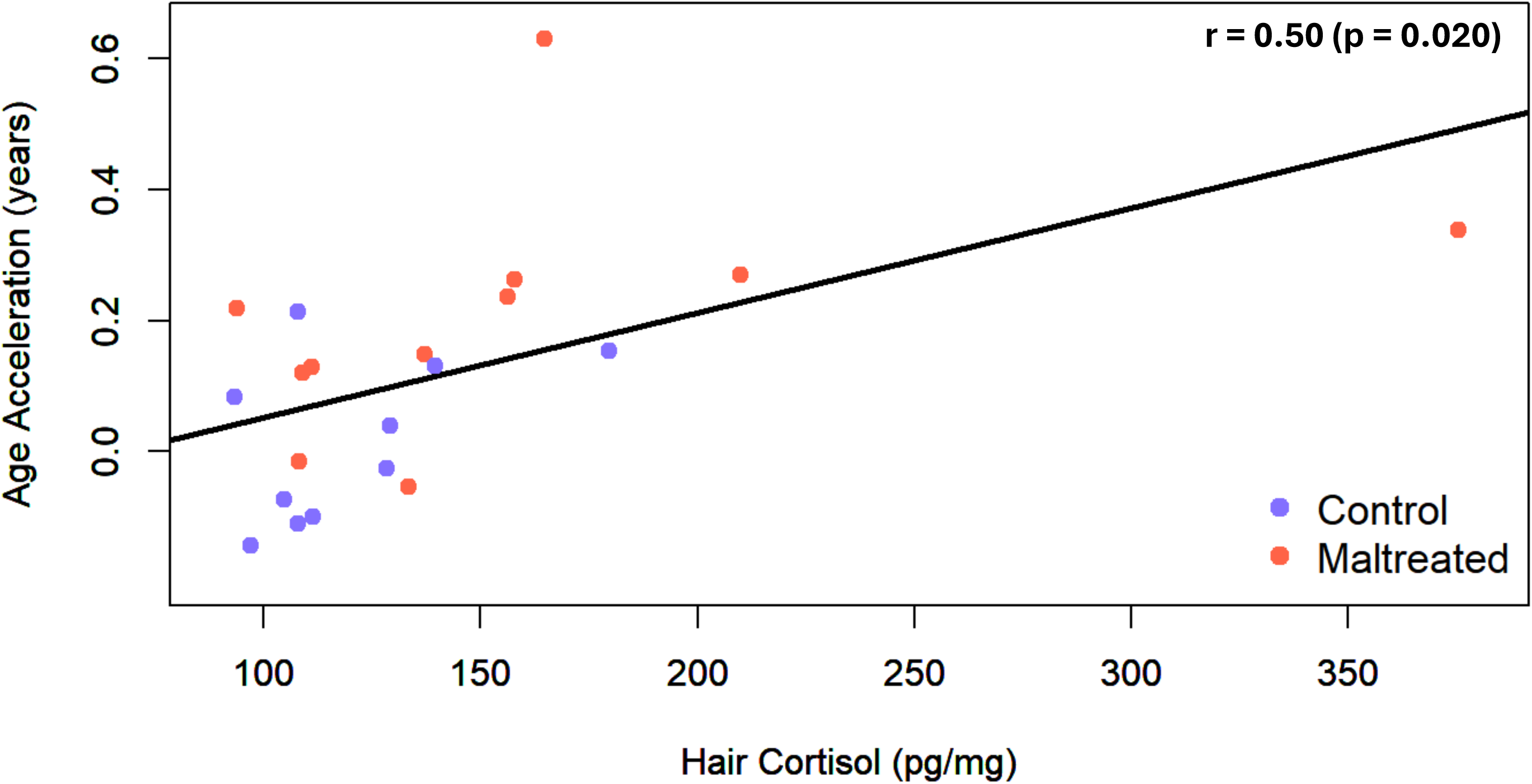
Age acceleration versus hair cortisol at age 0.5 years. Each maltreated and control individual is represented by a red or violet dot, respectively. The best fit line was fit to both maltreated and control individuals and has a slope of 0.0016 years/(pg/mg) – see *Methods, Modeling Age Acceleration* for details of the fitting. The Pearson correlation of age acceleration and hair cortisol is 0.50 with a p-value of 0.020 indicating a statistically significant correlation.

### Identification of DNA Methylation Loci Correlated with Age in the Rhesus Macaque Genome

When fitting the ECRM and ECYRM models, elastic net regularization selected 344 and 453 CpG sites, respectively, to include in the models. Because not all CpG sites are FDR-statistically significantly correlated with age, we computed the Pearson correlation of age with methylation for each of the selected CpGs. These correlations are displayed in **Tables S2 and S3** for ECRM and ECYRM, respectively, along with the corresponding FDR-adjusted p-values. A total of 269 CpGs showed an FDR-significant correlation with age in ECRM, while 373 CpGs showed an FDR-significant correlation with age in ECYRM. It is conceivable that the CpGs that play the largest role in aging are those that undergo the largest change in methylation with age. Therefore, in addition to the Pearson correlation, **Tables S2 and S3** show the rate of change of methylation with age (i.e. the rate of change of the methylation beta value).

**Tables S2 and S3** also display the closest gene to each CpG. We compared these genes to those of Horvath’s human multi-tissue clock.^2^ The following genes are shared by ECRM and the human clock: *ZBTB16*, *HOXB8, CBX7, TIMM17A, FXN,* and *SDC2*. The following genes are shared by ECYRM and the human clock: *ZBTB16, HOXB8, CBX7,* and *BIRC2*. While the methylation arrays differ between our study and Horvath’s such that we do not measure all the same CpGs, we do find the CpG cg05675373 is shared by ECYRM and the human clock, though none are shared by ECRM and the human clock. We also compared our clocks to a blood EWAS performed across numerous mammalian species^26^ that identified the top 1000 CpGs correlated with age and top 1000 CpGs anticorrelated with age and the genes closest to these CpGs. The following genes are shared by ECRM and the pan-mammalian EWAS: *CBLN1, GREM1, ZBTB16, SRSF7, RPN1, GABRB2, SLC1A3, MTDH, RIC3, SLC1A3, FOXP1*. The following genes are shared by ECYRM and the pan-mammalian EWAS: *FBXL7, ADGRL3, TUSC1, CA10, NOVA1, HOXD12, EPHA4, ZBTB16, GREM1, NR2E1, MIR548G, FOXP1, TYW3, BRINP3, HNRNPA1, CCAR1*. The CpG cg10501210 is shared by ECRM and the pan-mammalian EWAS, and cg10501210 and cg05675373 are shared by ECYRM and the pan-mammalian EWAS.

## Discussion

We created two epigenetic age estimators for rhesus macaques utilizing a large cohort of individuals. Our clocks achieve very high accuracy and are generally applicable as they are trained on animals from three different primate centers. Notably, the clocks achieve high accuracy for young individuals during critical developmental periods of life, which has not been demonstrated by rhesus macaque clocks previously.

Prior work by Horvath et al.^18^ and Goldman et al.^19^ developed epigenetic clocks for rhesus macaques using array- and sequencing-based methylation data. Horvath’s group developed the first rhesus-specific clock using a custom methylation array in n=281 samples from animals between 1.8 and 42 years of age. Their pan-tissue clock derived from blood, skin, fat, brain, kidney, liver, lung, and skeletal muscle tissue showed a Pearson correlation with chronological age of r=0.95 with a median absolute error (MAE) of 1.39 years while a separate blood-based clock (n=199) showed a correlation of r=0.95 and a MAE of 1.57 years.^18^ Additional pan-mammalian^26^ and pan-primate^27^ clocks derived from species including rhesus macaques provided insight into the biological aging across species. However, to the best of our knowledge, studies using these clocks have not provided evidence for an association between biological or environmental factors and epigenetic age acceleration in rhesus macaques. Goldman et al. used a high-throughput, sequencing-based method to profile DNA methylation in 493 free-ranging rhesus macaques between 1.44 and 28.82 years of age yielding a clock termed *RheMacAge* with a Pearson correlation of r=0.9 and MAE=1.42 years. The authors tested the association of epigenetic age acceleration with dominance rank; however, in contrast to a study in baboons,^20^ no association was detected. In addition, they found no association between epigenetic age acceleration and exposure to a natural disaster (Hurricane Maria).^19^

Utilizing the entire dataset of 1,008 samples, encompassing animals between 2 days and 23.4 years old, our *Epigenetic Clock for Rhesus Macaques* (ECRM) extends work by Horvath et al. and Goldman et al. to younger ages, and achieves a Pearson correlation with chronological age of r=0.983 and an MAE=0.210 years. We also specifically created an age estimator for younger animals including subjects between 2 days and 10 years of age (n=963) called the *Epigenetic Clock for Young Rhesus Macaques* (ECYRM), which achieves a Pearson correlation with chronological age of r=0.974 and an MAE of 0.148 years.

Like ours, prior studies have utilized human Illumina Infinium arrays to profile DNA methylation in rhesus macaques by leveraging the genetic similarity between humans and rhesus macaques.^28–31^ Although this choice may have weaknesses including the requirement for stringent QC and filtering and a limitation on DNA methylation sites that can be investigated, the cost, reliability, and robustness of the established array platforms and the ability to directly compare human and non-human primate data is an advantage. However, species other than rhesus may require sequencing-based technologies due to limited sequence overlap with human array platforms.

Age estimators are often used to investigate the effects of environmental and disease conditions on aging, morbidity, and mortality. Like other studies in humans, we applied our clock to a rhesus macaque model of ELA, namely infant maltreatment characterized by abusive behavior and neglect by the maltreating macaque mother towards the infant, causing pain and distress.^14–17,32^ Infant maltreatment in rhesus macaques is an aberrant behavior that can be observed in captivity and the wild occurring primarily in the first six months of life, when the infant is most dependent on the mother. Infant maltreatment has been linked to adverse developmental outcomes including the dysregulation of the hypothalamic-pituitary-adrenal (HPA) axis and disrupted development of brain circuits involved in emotion regulation.^17^ In fact, maltreated infants show elevated hair cortisol levels at 6 months of age indicating that maltreatment leads to a chronic activation of the HPA axis during the ELA experience.^17^ Prior evidence supports a causal relationship between HPA signaling and epigenetic aging. In a clinical study, Zannas et al. found dynamic DNA methylation changes in about 30% of the CpG sites of the Horvath human clock in response to dexamethasone, a synthetic agonist of the glucocorticoid receptor, suggesting that stress exposure via glucocorticoid signaling may influence aging trajectories.^33^

Using a case-control design, our data suggest that macaques experiencing ELA show an epigenetic age acceleration of 0.15 years (95% CI: 0.0064 to 0.29) compared to controls at 6 months of age, at the end of the main period of exposure to maltreatment. However, the wide 95% CIs indicate an imprecise determination of the degree to which maltreatment affects epigenetic aging. In fact, the effect size of age acceleration may be influenced by maltreatment severity, social buffering, or other stressors in the macaques’ environment. Therefore, we investigated hair cortisol levels as a more precise measure of chronic stress exposure. As expected, we found a precisely determined direct correlation between hair cortisol accumulation and epigenetic age acceleration in 6-month-old macaques. Individuals that differ in hair cortisol by two standard deviations differ in age acceleration by 0.20 years (95% CI of 0.054 to 0.34 years). However, we did not find sufficient evidence for age acceleration prior to 6 months, nor do we have evidence that age acceleration persists at 2 years of age. A large body of literature suggests long-lasting effects of childhood maltreatment, trauma exposure and stress in general on epigenetic aging in humans.^25^ However, cumulative effects of chronic stress exposure and recency/concurrent stress effects may be harder to disentangle in human studies.

In conclusion we developed two epigenetic age estimators for rhesus macaques and provided evidence for accelerated epigenetic aging in response to infant maltreatment, which is likely driven by chronic HPA axis activation. Future studies are required to provide evidence for the applicability of these clocks given that both are trained on chronological age but not trained on other markers of morbidity and mortality. Nevertheless, with their high degree of accuracy, our epigenetic clocks may be applicable in studies on aging and age-related disorders in rhesus macaques, a species highly relevant for translational aging research.

## Methods

### Cohort Information

#### Wisconsin National Primate Research Center (WNPRC)

Cross-sectional blood samples were collected as part of a large research program on early life in rhesus macaques, including animals assessed for anxious temperament.^34,35^ A total of 516 samples were included in the analyses. A subset of 243 samples were derived from the WNPRC 1 cohort, while 273 samples were derived from the Harlow Center for Biological Psychology (WNPRC 2 cohort). This research study was approved by the Institutional Animal Care and Use Committee of the University of Wisconsin (protocol # G005809). Chronological age at the time of the blood draw was determined by utilizing pedigree records.

#### Emory National Primate Research Center (ENPRC)

We utilized whole blood samples from two independent cohorts described before. Briefly, ENPRC 1 is a cohort of 42 individuals with 192 samples from a longitudinal, developmental, study of ELA in the form of infant maltreatment from birth through the juvenile period.^14,17,36,37^ Peripheral blood samples were collected as described below at day 2 after birth, week 2, month 1, month 3, month 6, and 1.8-2.2 years of age. This research study was approved by the Institutional Animal Care and Use Committee of Emory University (protocol #2001377). Blood samples for the ENPRC 2 cohort were collected from birth through puberty, including 44 individuals and 212 samples, as part of a study on the developmental effects of social subordination stress and exposure to an obesogenic diet compared to a standard diet. This research study was approved by the Institutional Animal Care and Use Committee of Emory University (protocol #2003417). Chronological age at the time of the blood draw was determined by utilizing ENPRC records.

#### Caribbean Primate Research Center (CPRC)

Peripheral blood samples from a cross-sectional study on fear in 88 free-ranging individuals from Cayo Santiago were collected as described before.^31^ Chronological age at the time of the blood draw was determined by utilizing Center records. This research study was approved by the Institutional Animal Care and Use Committee of the University of Puerto Rico (protocol #A3340109).

### Sample Preparation and Processing

Whole blood samples were drawn in EDTA tubes and frozen until further analysis. The Gentra Puregene Blood Kit (Qiagen) was used to extract genomic DNA at McLean Hospital or AKESOgen Inc (Peachtree Corners, GA), where DNA samples were also further processed and analyzed. For the WNPRC/Harlow cohort, the buffy coat layer obtained from whole blood was shipped to the Baylor College of Medicine for DNA purification using the Gentra Puregene Blood Kit or phenol/chloroform extraction. These DNA samples were then shipped to McLean Hospital for analysis. PicoGreen was used to measure DNA concentrations. Bisulfite conversion was performed using the EZ-96 DNA methylation kit (Zymo). DNA was then hybridized to human Illumina Infinium MethylationEPIC v1.0 microarrays and scanned on an iScan (Illumina). Illumina methylation assays quantify DNA methylation levels with fluorescence intensity of the methylated and un-methylated signals, from which the β values can be calculated.^38^ The β-value for a particular CpG site is the fraction of copies of that CpG that are methylated. Processing of DNA methylation data is described below.

### Hair Cortisol

Hair cortisol concentrations from the ENPRC 1 cohort were measured to examine chronic hypothalamic-pituitary-adrenal (HPA) axis activation due to the infant maltreatment ELA as described before.^17^ Briefly, hair was shaved from the back of the neck to measure cortisol accumulation in the growing hair shaft. Hair samples were processed and assayed following published procedures,^39^ and cortisol was measured using enzyme immunoassay (#1-3002, Salimetrics, Carlsbad, CA). Intra- and inter-assay coefficients of variation were <10%.

### DNA Methylation Data Processing

DNA methylation data processing was performed in R (version 4.1.2) and RStudio (version 2022.12.0+353.pro20). Using the wateRmelon package (version 2.0.0),^40^ we carried out the following steps to filter the samples and CpG probes: Only probes that mapped to the macaque genome were included.^31^ Bisulfite conversion rates were computed with the *bscon* function, and we only included samples in our analysis if they achieved a bisulfite conversion rate of greater than 80%. Next, detection p-values were computed using the *pfilter* function, and samples were excluded if greater than 5% of their CpG sites had a detection p-value greater than 0.05. Also using *pfilter*, CpG sites were excluded if either (1) the detection p-value was greater than 0.05 in greater than 5% of samples, or (2) the bead count was less than 3 in greater than 5% of samples. Next, we excluded outlier samples that were detected through a principal component analysis using the *outlyx* function. Then, probes targeting the X or Y chromosomes were excluded. Next, the *dupicc* function from the ENmix package (version 1.30.3)^41^ was applied to 12 pairs of replicate samples to assess the intraclass correlation coefficient (ICC) for each CpG. Based on these ICCs, we excluded from our analysis all CpG sites whose ICCs were not statistically significantly greater than 0 (i.e. p-value cut-off of 0.05). After filtering, we were left with 1,008 samples (out of 1,136 total samples) and 194,251 CpG sites for inclusion in the epigenetic clock model (of which 344 or 453 sites were selected by elastic net regularization for ECRM and ECYRM, respectively, discussed below). Finally, the data was normalized using quantile normalization followed by beta-mixture quantile normalization (BMIQ),^42^ and it is this normalized data that was then used to create the epigenetic clock model.

### Epigenetic Age Estimator Model

The epigenetic clock models are multiple linear models, where the DNA methylation levels (i.e. normalized methylation beta values) are the independent variables and the rhesus macaque’s age is the dependent variable. The models are fit with elastic net regularization, performed by the *glmnet* function in the glmnet R package (version 4.1.7)^43^ using the Gaussian family objective function. The elastic net hyperparameters are λ, which controls the strength of the elastic net penalty, and α, the elastic net mixing parameter.

The epigenetic clock models are given by

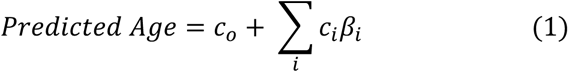

where *β_i_* is the normalized methylation beta value for the *i^th^* CpG, and *c_i_* is the corresponding coefficient that was determined using glmnet (for the coefficient values, see “Clock Coefficient” in **Tables S2 and S3**). The summation is performed over all CpGs chosen by the elastic net regularization (344 and 453 CpGs for ECRM and ECYRM, respectively). *c_0_* is the intercept parameter found by glmnet, which is 10.696 years for ECRM and 4.768 years for ECYRM.

We created two clocks that are available in the R language as tools for other researchers to use. (1) We created the ECRM model, which is available at https://github.com/klengellab/MonkeyAge. We performed 5-fold cross-validation on all 1,008 samples to test hyperparameter sets using a grid search across the range of all possible values. We found that α = 0.2 and λ = 0.174 are the hyperparameter values that give the maximum Pearson correlation between the true age and predicted age. Using this hyperparameter set and all 1008 samples, we retrained the model and obtained the resulting model coefficients (provided in **Supplementary File 1 and Table S2**). (2) We created the ECYRM model in the same fashion, using only samples below the age of 10 years (n = 963). We found that α = 0.2 and λ = 0.199 are the hyperparameter values that give the maximum Pearson correlation between the true age and predicted age, as well as minimum RMSD. ECYRM is available at https://github.com/klengellab/MonkeyAge **and Table S3**. For ECYRM, a cut-off of 10 years for the training data was chosen because including individuals older than 10 (as in ECRM) results in less accuracy for ages under 6 years, as discussed in *Results*. A cut-off of 10 rather than 6 years was used because of a systematic error in the epigenetic clock model. Specifically, it underestimates the ages of individuals whose true ages are near the upper end of the age range of the training data. We found that setting the cut-off to 6 years resulted in macaques aged 5 to have predicted ages that are too low. Therefore, we included individuals up to 10 years in the training, avoiding the systematic error present at age 5. The systematic error is, however, still present for individuals over the age of 6 years, as can be seen in **Figure 2**. Therefore, we only apply ECYRM to study macaques under the age of 6 years.

We determined the accuracy of the two clocks by performing nested cross-validation. (1) To estimate the accuracy of the ECRM model, we performed nested cross-validation on all 1,008 samples. The inner loop of 9-fold cross-validation served to optimize the hyperparameters for elastic net regularization such that the Pearson correlation between the true age and predicted age was maximized. The outer loop of 10-fold cross-validation made use of these optimized hyperparameters and trained the model on 90% of the samples and then predicted the ages of the other 10% of samples. (2) We determined the accuracy of the ECYRM model in the same fashion but using only the 963 samples below the age of 10 years.

We created a third epigenetic clock for use in our study of infant maltreatment, which we call ECYRM_modified_. The clock uses 771 blood samples – these are all the samples below the age of 10 years, excluding all individuals in the ENPRC 1 cohort. After performing 5-fold cross-validation to select hyperparameters, we fit the model on all 771 blood samples and then used this model to predict the ages of the individuals in the ENPRC 1 cohort, which served as an independent test set. The epigenetic age accelerations of the ENPRC 1 cohort were then further analyzed in the context of maltreatment and cortisol, as described in *Results*.

We note that not all samples contain measurements for all CpG sites. We use mean imputation to solve this issue – i.e. whenever a CpG measurement is missing in a sample in the training or validation set, its value is taken to be the mean value of that CpG across all the samples in the training set that have measurements of that CpG. If a CpG measurement is missing in the testing data, we use the mean value of that CpG in the training and validation data. Of all of our data fed into the elastic net regularization algorithm, only a small percentage (0.79%) of the methylation beta values were imputed, and of the data that was used to train the clocks (i.e. the CpGs selected by the elastic net), 6.9% and 5.3% of methylation beta values were imputed for ECRM and ECYRM, respectively.

### Age Acceleration Outlier Detection

To prevent outliers from obscuring the effects of maltreatment in the ENPRC 1 cohort, we evaluated whether any sample in this cohort was an outlier in terms of its age acceleration. Outliers were determined using Grubbs’ test with a p-value cut-off of 0.01 (cut-off chosen to err on the side of not removing data from our analyses), comparing the age accelerations of individuals at the same time point within the same group (i.e. the control group or the maltreatment group). We identified one outlier to remove from our age acceleration analysis (Grubb’s test p-value of 0.00020). This was a control macaque at age 0.5 years whose age acceleration was 0.87 years, while the 10 other control macaques at this time point had an age acceleration between -0.14 and 0.21 years.

We also investigated if any individuals did not follow the same relationship between age acceleration and hair cortisol as the other rhesus macaques. Using the *outlierTest* function from the “car” R package,^44^ we performed a Bonferroni outlier test to determine whether any macaques had a statistically significant studentized residual when performing a linear regression of age acceleration on hair cortisol and sex at 0.5 years. No macaques reached a significance threshold of Bonferroni adjusted p-value < 0.01, so we did not remove any individuals from the analysis on account of this test.

### Modeling Age Acceleration

As shown in equation (2), we investigated the effect of maltreatment by applying a multiple linear model for age acceleration for the ENPRC 1 cohort at each time point:

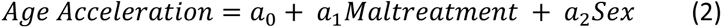

The variables are as follows: *Maltreatment* is 1 if the macaque mother maltreated its infant and 0 if the mother did not (i.e. she was a competent caregiver). Sex is 1 if the macaque was female and 0 if male. The parameters *a*_1_ and *a*_2_ are the effects of each variable on age acceleration. *a*_1_ is the change in age acceleration due to maltreatment: the values for each time point are displayed in **Table 1**. *a*_2_ is the change in age acceleration associated with the macaque’s sex, which we find to be -0.0029 years (95% CI: -0.14 to 0.14) at the 0.50-year time point. Similarly, the 95% CIs overlap 0 at all other time points, except at 1.8-2.2 years when *a*_2_ = -0.28 (95% CI: -0.52 to -0.044). 2_0_ is the intercept term, included in the model in case there is a systematic error in the epigenetic clock’s ability to predict ages in the independent test cohort, though we find *a*_0_ to be minimal.

As shown in equation (3), we investigated the correlation of cortisol and age acceleration by applying a multiple linear model for the ENPRC 1 cohort at 6 months:

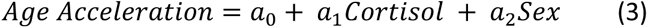

*Cortisol* is the hair cortisol measurement divided by the standard deviation of hair cortisol across this cohort at 0.5 years of age. (We did not include *Maltreatment* and *Cortisol* in the same model because they are approximately colinear). *a*_1_ is the change in age acceleration associated with an increase in hair cortisol of one standard deviation (the standard deviation of hair cortisol levels at this time in the ENPRC 1 cohort is 62 pg/mg). We find through Bayesian parameter estimation that the value of *a*_2_ is 0.10 years (95% CI of 0.027 to 0.17), as discussed in *Results*. The value of *a*_2_ is -0.059 years (95% CI of - 0.20 to 0.088). The relationship between hair cortisol and age acceleration is depicted graphically in **Figure 3**, where the best fit line is given by y = 0.0016x - 0.11. Here the slope of 0.0016 is equivalent to *a*_1_ but is calculated without standardizing the hair cortisol by the standard deviation of hair cortisol. The intercept of -0.11 represents *a*_0_ plus the mean effect of the other covariate: namely, the intercept is given by *a*_0_ + *f*_2_*a*_2_, where *f*_2_ is the fraction of individuals that are female.

To estimate the values of the model parameters and their 95% CIs, we perform Bayesian parameter estimation. To do so, we employ an error model that is a Gaussian distribution with mean 0 and standard deviation parameter σ. (While equation 2 represents the expected age acceleration for a macaque, the error model represents the fact that there is unexplained variation that makes an individual’s age acceleration deviate from the expected value). Therefore, the likelihood function that we use in Bayesian parameter estimation is a Gaussian with standard deviation σ and a mean given by the right-hand side of equation 2.

As we had no prior knowledge about the values of these parameters, we employed a non-informative prior probability distribution for the parameters, specifically the Jeffreys prior. Because our likelihood function is a linear model with a Gaussian error model, the Jeffreys prior is the following: For *a_i_*, the prior is a uniform distribution, where -10^6^ < *a_i_* < 10^6^; for σ, the prior probability density function is a Pareto distribution, given by *PDF*(σ) = (*M* + 1)*c^M^*^+1^σ^(*M*+2)^, where σ > c. c = 10^-6^ and M is the number of independent variables, which is 2 in the cases of equations 2 and 3. (These bounds on *a_i_* and σ were chosen to be quite wide to emulate a truly noninformative prior). The Bayesian parameter estimation was carried out using the Gibbs sampling program JAGS^45^ version 4.3.1 via the R package rjags^46^ until simulation convergence was reached (10^4^ trials for burn-in followed by 7*10^5^ simulation trials).

Epigenetic clocks are trained using the hyperparameter set that optimizes a particular performance metric, such as Pearson correlation between true age and predicted age, median absolute error, mean absolute error, or RMSD. However, one must make an arbitrary choice when deciding which performance metric the clock should be optimized for. Using hyperparameter sets that optimize different performance metrics will often yield slightly different age predictions. To verify the robustness of our conclusions regarding infant maltreatment and cortisol’s correlation with age acceleration, we fit the epigenetic clock four times, each time optimizing based on a different metric (Pearson correlation, median absolute error, mean absolute error, or RMSD). We then used the resulting age accelerations in equations 2 and 3. We found that regardless of the performance metric used for optimization, the mean effect size at 6 months for hair cortisol is always between 0.081 and 0.10 years and for maltreatment is always between 0.11 and 0.15 years, and there is always ≥ 0.95 probability that these effect sizes are positive. Note that all model coefficients reported in this manuscript were calculated when using the Pearson correlation as the performance metric of choice.

## Data Availability

The analysis code is publicly available at https://github.com/klengellab/MonkeyAge. Relevant data has been deposited at GEO under the accession number GSE272604.

## Acknowledgements

We want to thank Drs. Jeffrey Rogers and Muthuswamy Raveedran (Baylor College of Medicine, Houston, TX) for help in providing the DNA samples for the WNPRC. The ENPRC work was supported by funding from the National Institutes of Health (NIH) grant numbers MH078105, MH086203, HD077623, HD088931, HD097524, AG070704, the NIH’s Office of the Director, Office of Research Infrastructure Programs, P51OD011132 (ENPRC Base grant) and S10OD010757–01 (ENPRC Biomarkers Core, also supported by the ENPRC Base Grant). The authors want to thank Anne Glenn, Christine Marsteller, Dora Guzman, Natalie Brutto, Kelly Bailey, Trina Jonesteller, Sara Dicker, Jodi Godfrey and the staff at the ENPRC for the excellent technical support and animal care provided during the studies. The ENPRC is fully accredited by AAALAC, International. TK was supported by HD102974, HD088931, HD097524, AG070704, P50MH115874, the Connor Group Kids and Community Partners, and ERA-NET Neuron 01EW2003.

## Supplementary Information

**Table S1.** Number of Samples for Each Age Used in the Epigenetic Clock Models. Displayed are the number of blood samples from rhesus macaques of each age group in each cohort. The ECRM model uses all samples, while the ECYRM model uses all samples except ages 10-23 years old (y.o.). We use data from five cohorts of rhesus macaques: two cohorts from the Emory National Primate Research Center (ENPRC), two from the Wisconsin National Primate Research Center (WNPRC), and one from the Caribbean Primate Research Center (CPRC). In cohorts CPRC and WNPRC 1 and 2, a single blood sample was taken from each individual. However, in ENPRC 1 and 2, multiple blood samples were taken from each individual, with each sample taken at a different time point.

**Table S2.** List of CpGs in the ECRM Model, Ordered by Absolute Value of Pearson Correlation. Displayed in the 2^nd^ column is the nearest gene to each CpG. Shown in the 3^rd^ column is the distance between the CpG and the nearest gene. A distance of 0 bp indicates that the CpG is located within the coding region of the gene; a negative distance indicates the CpG is upstream of the coding region, while a positive distance indicates the CpG is downstream of the coding region. The 4^th^ column shows the Pearson correlation between age and the CpG’s methylation beta value, and the 5^th^ column is the FDR-adjusted p-value for the Pearson correlation. The 6^th^ column is the rate of change of the methylation beta value (units of 1/year) and the 7^th^ column is the FDR-adjusted p-value testing whether the rate of change is significantly different from 0. The 8^th^ column is the coefficient of each CpG in the epigenetic clock model.

**Table S3.** List of CpGs in the ECYRM Model, Ordered by Absolute Value of Pearson Correlation. See the legend of Table S2 for details.

## References

1. Hannum, G. et al. Genome-wide methylation profiles reveal quantitative views of human aging rates. Mol. Cell 49, 359–367 (2013).

2. Horvath, S. DNA methylation age of human tissues and cell types. Genome Biol. 14, 3156 (2013).

3. Perna, L. et al. Epigenetic age acceleration predicts cancer, cardiovascular, and all-cause mortality in a German case cohort. Clin. Epigenetics 8, 64 (2016).

4. Marioni, R. E. et al. DNA methylation age of blood predicts all-cause mortality in later life. Genome Biol. 16, 25 (2015).

5. Marioni, R. E. et al. The epigenetic clock is correlated with physical and cognitive fitness in the Lothian Birth Cohort 1936. Int. J. Epidemiol. 44, 1388–1396 (2015).

6. Levine, M. E., Lu, A. T., Bennett, D. A. & Horvath, S. Epigenetic age of the pre-frontal cortex is associated with neuritic plaques, amyloid load, and Alzheimer’s disease related cognitive functioning. Aging 7, 1198–1211 (2015).

7. Binder, A. M. et al. Faster ticking rate of the epigenetic clock is associated with faster pubertal development in girls. Epigenetics 13, 85–94 (2018).

8. Peng, C. et al. Epigenetic age acceleration is associated with allergy and asthma in children in Project Viva. J. Allergy Clin. Immunol. 143, 2263–2270.e14 (2019).

9. Davis, E. G. et al. Accelerated DNA methylation age in adolescent girls: associations with elevated diurnal cortisol and reduced hippocampal volume. Transl. Psychiatry 7, e1223–e1223 (2017).

10. Boks, M. P. et al. Longitudinal changes of telomere length and epigenetic age related to traumatic stress and post-traumatic stress disorder. Psychoneuroendocrinology 51, 506–512 (2015).

11. Jovanovic, T. et al. Exposure to Violence Accelerates Epigenetic Aging in Children. Sci. Rep. 7, 8962 (2017).

12. Sumner, J. A., Colich, N. L., Uddin, M., Armstrong, D. & McLaughlin, K. A. Early Experiences of Threat, but not Deprivation, Are Associated With Accelerated Biological Aging in Children and Adolescents. Biol. Psychiatry 85, 268–278 (2019).

13. Phillips, K. A. et al. Why primate models matter. Am. J. Primatol. 76, 801–827 (2014).

14. Drury, S. S. et al. Shaping long-term primate development: Telomere length trajectory as an indicator of early maternal maltreatment and predictor of future physiologic regulation. Dev. Psychopathol. 29, 1539–1551 (2017).

15. Howell, B. R. & Sanchez, M. M. Understanding behavioral effects of early life stress using the reactive scope and allostatic load models. Dev. Psychopathol. 23, 1001–1016 (2011).

16. Maestripieri, D. The biology of human parenting: insights from nonhuman primates. Neurosci. Biobehav. Rev. 23, 411–422 (1999).

17. McCormack, K. M. et al. The developmental consequences of early adverse care on infant macaques: A cross-fostering study. Psychoneuroendocrinology 146, 105947 (2022).

18. Horvath, S. et al. Epigenetic clock and methylation studies in the rhesus macaque. GeroScience 43, 2441–2453 (2021).

19. Goldman, E. A. et al. A generalizable epigenetic clock captures aging in two nonhuman primates. 2022.11.01.514719 Preprint at 10.1101/2022.11.01.514719 (2022).

20. Anderson, J. A. et al. High social status males experience accelerated epigenetic aging in wild baboons. eLife 10, e66128 (2021).

21. Sugden, K. et al. Patterns of Reliability: Assessing the Reproducibility and Integrity of DNA Methylation Measurement. Patterns 1, 100014 (2020).

22. Mann, D. R., Akinbami, M. A., Gould, K. G., Tanner, J. M. & Wallen, K. Neonatal treatment of male monkeys with a gonadotropin-releasing hormone agonist alters differentiation of central nervous system centers that regulate sexual and skeletal development. J. Clin. Endocrinol. Metab. 76, 1319–1324 (1993).

23. Catchpole, H. R. & Van Wagenen, G. Physical growth of the Rhesus monkey (Macaca mulatta). Am. J. Phys. Anthropol. 14, 245–273 (1956).

24. Wilson, M. E. & Tanner, J. M. Long-term effects of recombinant human growth hormone treatment on skeletal maturation and growth in female rhesus monkeys with normal pituitary function. J. Endocrinol. 130, 435–441 (1991).

25. Musci, R. J. et al. Using Epigenetic Clocks to Characterize Biological Aging in Studies of Children and Childhood Exposures: a Systematic Review. Prev. Sci. Off. J. Soc. Prev. Res. 24, 1398–1423 (2023).

26. Lu, A. T. et al. Universal DNA methylation age across mammalian tissues. *Nat*. Aging 3, 1144– 1166 (2023).

27. Horvath, S. et al. Pan-primate studies of age and sex. GeroScience 45, 3187–3209 (2023).

28. Pichon, F. et al. Analysis and Annotation of DNA Methylation in Two Nonhuman Primate Species Using the Infinium Human Methylation 450K and EPIC BeadChips. Epigenomics 13, 169–186 (2021).

29. Ong, M.-L. et al. Infinium Monkeys: Infinium 450K Array for the Cynomolgus macaque (Macaca fascicularis). G3 GenesGenomesGenetics 4, 1227–1234 (2014).

30. Housman, G., Quillen, E. E. & Stone, A. C. Intraspecific and interspecific investigations of skeletal DNA methylation and femur morphology in primates. Am. J. Phys. Anthropol. 173, 34–49 (2020).

31. Bravo-Rivera, H. et al. Innate fear responses are reflected in the blood epigenome of rhesus macaques. 2020.11.05.369538 Preprint at 10.1101/2020.11.05.369538 (2020).

32. McCormack, K., Sanchez, M. M., Bardi, M. & Maestripieri, D. Maternal care patterns and behavioral development of rhesus macaque abused infants in the first 6 months of life. Dev. Psychobiol. 48, 537–550 (2006).

33. Zannas, A. S. et al. Lifetime stress accelerates epigenetic aging in an urban, African American cohort: relevance of glucocorticoid signaling. Genome Biol. 16, 266 (2015).

34. Oler, J. A. et al. Amygdalar and hippocampal substrates of anxious temperament differ in their heritability. Nature 466, 864–868 (2010).

35. Fox, A. S. et al. Intergenerational neural mediators of early-life anxious temperament. Proc. Natl. Acad. Sci. 112, 9118–9122 (2015).

36. Howell, B. R. et al. Disentangling the effects of early caregiving experience and heritable factors on brain white matter development in rhesus monkeys. NeuroImage 197, 625–642 (2019).

37. Morin, E. L. et al. Developmental outcomes of early adverse care on amygdala functional connectivity in nonhuman primates. Dev. Psychopathol. 32, 1579–1596 (2020).

38. Sala, C. et al. Evaluation of pre-processing on the meta-analysis of DNA methylation data from the Illumina HumanMethylation450 BeadChip platform. PLOS ONE 15, e0229763 (2020).

39. Meyer, J., Novak, M., Hamel, A. & Rosenberg, K. Extraction and analysis of cortisol from human and monkey hair. J. Vis. Exp. JoVE e50882 (2014) doi:10.3791/50882.

40. Schalkwyk, L. C., et al. wateRmelon: Illumina 450 and EPIC methylation array normalization and metrics. Bioconductor version: Release (3.17) 10.18129/B9.bioc.wateRmelon (2023).

41. Xu, Z., Niu, L. & Taylor, J. ENmix: Quality control and analysis tools for Illumina DNA methylation BeadChip. Bioconductor version: Release (3.17) 10.18129/B9.bioc.ENmix (2023).

42. Wang, T. et al. A systematic study of normalization methods for Infinium 450K methylation data using whole-genome bisulfite sequencing data. Epigenetics 10, 662–669 (2015).

43. Friedman, J., et al. glmnet: Lasso and Elastic-Net Regularized Generalized Linear Models. (2023).

44. Fox J, Weisberg S. An R Companion to Applied Regression. (Sage, Thousand Oaks CA, 2019).

45. JAGS - Just Another Gibbs Sampler. https://mcmc-jags.sourceforge.io/.

46. Plummer, M., Stukalov, A. & Denwood, M. rjags: Bayesian Graphical Models using MCMC. (2023).

